# VarExp: Estimating variance explained by Genome-Wide GxE summary statistics

**DOI:** 10.1101/224634

**Authors:** Vincent Laville, Amy R. Bentley, Florian Privé, Xiafoeng Zhu, Jim Gauderman, Thomas W Winkler, Mike Province, DC Rao, Hugues Aschard, on behalf of the CHARGE Gene-Lifestyle Interactions Working Group

## Abstract

Many genomic analyses, such as genome-wide association studies (GWAS) or genome-wide screening for Gene-Environment (GxE) interactions have been performed to elucidate the underlying mechanisms of human traits and diseases. When the analyzed outcome is quantitative, the overall contribution of identified genetic variants to the outcome is often expressed as the percentage of phenotypic variance explained. In practice, this is commonly estimated using individual genotype data. However, using individual-level data faces practical and ethical challenges when the GWAS results are derived in large consortia through meta-analysis of results from multiple cohorts. In this work, we present a R package, “VarExp”, that allows for the estimation of the percentage of phenotypic variance explained by variants of interest using summary statistics only. Our package allows for a range of models to be evaluated, including marginal genetic effects, GxE interaction effects, and main genetic and interaction effects jointly. Its implementation integrates all recent methodological developments on the topic and does not need external data to be uploaded by users.

The R source code, tutorial and associated example are available at https://gitlab.pasteur.fr/statistical-genetics/VarExp.git.

## 1 Introduction

Many genome-wide association studies (GWAS) (Welter *et al*., 2014) or genome-wide screenings incorporating gene-environment (GxE) interactions (Aschard *et al*., 2012; McAllister *et al*., 2017) have been performed to better understand underlying mechanisms of human traits and diseases. When the analyzed outcome is continuous, a commonly-used measure to judge the overall impact of the significant associations is the percentage of phenotypic variance explained. A standard way of estimating this percentage is to compare the coefficients of determination between the model including both covariates and the significantly associated variants and/or interactions and the model with the covariates only. This estimation requires individual genotype and phenotype data which can be challenging in meta-analyses performed in big consortia as pooling data from multiple cohorts raises practical and ethical issues. However, an alternative is to use only GWAS or genome-wide GxE summary statistics to estimate the percentage of variance explained by new discoveries. Recently, several methods (Pare *et al*., 2016; Shi *et al*., 2016) have been developed to apply this strategy to marginal genetic effects while taking into account linkage disequilibrium (LD) between variants, and addressing statistical issues related to finite sample size and Single Nucleotide Polymorphisms (SNP) correlation matrices. Yet, application can remain challenging in practice, as the derivation of variance explained requires not only summary statistics but also external information on LD from a reference panel. More importantly, these works only focused on marginal genetic effects, while genome-wide GxE and joint effect GWAS are now commonly performed and face the same need. Here, we address this gap, extending the methodology to GxE screening and implementing in R package *VarExp* to rapidly and easily estimate the percentage explained by variants and/or interactions of interest using only meta-analysis summary statistics from GWAS.

## 2 Implementation

### 2.1 Percentage of variance explained by genetic and/or interaction effects

#### Marginal model

Consider a set of *K* SNPs (*G_k_*)_*k*=1…*K*_,coded additively as {0,1,2} and a quantitative outcome *Y*. The marginal genetic effect *α_G.k_* of SNP *G_k_* is estimated in the marginal model:

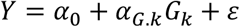

Shi et al. (Shi *et al*., 2016) proposed a first naïve estimator to derive the variance explained by genetic effects, *f_G_* using summary statistics:

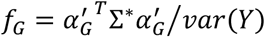

where 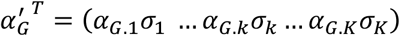, *σ_k_* denotes the standard deviation of SNP *G_k_*, Σ* is the Moore-Penrose generalized inverse of the genotype correlation matrix Σ. However, finite sample size implies statistical noise in both the effect sizes and the correlation matrix estimations which can induce bias in the estimation of *f_G_*. Shi et al. (Shi *et al*., 2016) derived a general formula that addresses this issue:

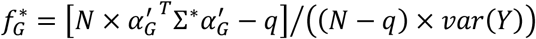

where *N* and *q* denote respectively the sample size and the rank of the correlation matrix.

#### Model with GxE interaction

Now consider an exposure *E* (either binary or quantitative). The main effect *α_G.k_* of the SNP *k* and the interaction effect *α_INT.k_* can be estimated using a single-SNP model with an interaction term:

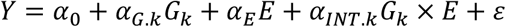

We show in Supplementary file that, when re-parameterizing the effect estimates of the above model to obtain parameters from a fully standardized model, the percentage of variance explained by interactions effects *f_I_* or jointly by genetic and interaction effects *f_G+I_* can also be derived using summary statistics only:

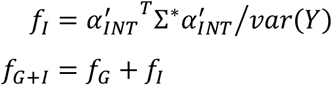

where 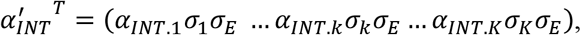, *σ_k_* and *σ_E_* are respectively the standard deviation of SNP *k* and *E*. Note that in this model, *f_G_* is computed using effect sizes from the interaction model. However, for the reasons discussed above, we define our final estimators, 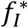 and 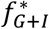, by applying the same corrections as proposed for the *f_G_* estimator by Shi et al (see 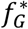 equation).

### 2.2 Estimating the genotype correlation matrix

As shown in section 2.1, the derivation requires the genotype correlation matrix. When this is not available from the data, this correlation matrix can be estimated using genotype data of founders in the appropriate ancestry from a reference panel such as the 1000 Genomes (Abecasis *et al*., 2013). In our R package, we implemented a transparent function that derives this correlation matrix from curated 1000 Genomes Phase 3 VCF files generated as part of another project (http://bochet.gcc.biostat.washington.edu/beagle/1000Genomesphase3v5a/) either through a web access (for small number of SNPs) or from local data files (for larger number of SNPs, see Supplementary File and Supplementary Figure 2). To avoid matrix inversion issues, we also implemented an option to remove SNPs with perfect correlation of 1 with another SNP in the matrix.

### 2.3 Input files

The user needs to provide two mandatory input files. The first input file contains the meta-analysis summary statistics with several mandatory columns: the chromosome and physical position (currently NCBI Build 37) of the variants, the tested allele and its frequency, the estimated genetic, interaction or both effect(s). The second file gives the sample size, the mean and variance for both the studied outcome and the exposure in each cohort included in the meta-analyses (or a subset) when considering interaction effects. If the outcome is binary, mean and variance can be replaced by counts of exposed individuals and sample sizes per cohort.

### 2.4 Application example

In practice, application is performed in 3 main steps: 1) after loading the two mandatory input files described in section 2.4, users call the *getGenoCorMatrix()* function to estimate the SNP correlation matrix as described in section 2.2. 2) parameters (mean and variance) of both the outcome and the exposure in the pooled sample are then computed using the functions *calculateParamsFromlndParams()* or *calculateParamsFromCounts()*, respectively. 3) finally, the *calculateVarExp()* function estimates the percentage of phenotypic variance explained by main genetic effects and/or interaction effects. To illustrate the performances of our package, we performed a simulation study (see Supplementary File for details of the framework) comparing the adjusted coefficients of determination from regressions and the estimates obtained using *VarExp* across 1,000 replicates. Figure 1 and Supplementary Figure 1 demonstrate the high accuracy of our estimator with an intraclass correlation coefficient between the coefficients of determination and their estimations equal to 0.99, 0.98 and 0.99 for the marginal genetic effects, interaction effects and joint effects respectively.

**Figure 1.**
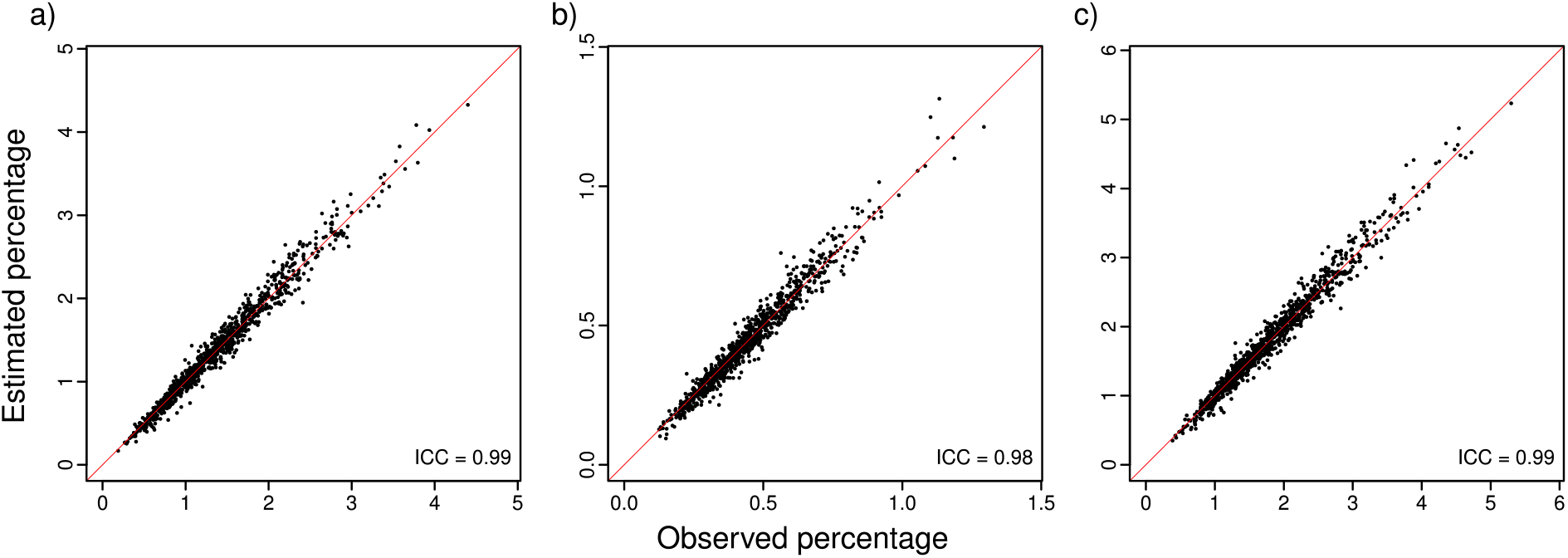
Summary statistics based estimations of the percentage of phenotypic variance explained by (a) main genetic (b) interaction and (c) joint effects, against regression coefficients of determination derived in individual-level data. The red line corresponds to y = x and *ICC* is the intraclass correlation coefficient between the two series.

## 3 Concluding remarks

In this work, we provide a simple pipeline to estimate the percentage of phenotypic variance explained by genetic effects, GxE interaction effects or their joint contribution using summary statistics only. Our approach makes this estimation straightforward in large-scale consortia where pooling individual genotype data can be extremely challenging. The approach is implemented in the R package *VarExp*.

## Acknowledgements

We gratefully acknowledge all contributors to the CHARGE Gene-Lifestyle Interactions Working Group.

